# Light-inducible proximity labelling *in vivo* captures sex-specific RNA at excitatory synapses

**DOI:** 10.64898/2026.01.28.702434

**Authors:** Joshua W. A. Davies, Valeria Suhinin, Arie Brueckner, Gabriel John A. Araneta, Laura J. Leighton, Alexander D. Walsh, Sachithrani U. Madugalle, Timothy Young, Hao Gong, Mason R. B. Musgrove, Haobin Ren, Marcin Kielar, John Yu-luen Lin, Ying Li, Timothy W. Bredy, Paul R. Marshall

## Abstract

Local RNA regulation is essential for synaptic plasticity, yet the full repertoire of RNA species and associated isoforms within specific synaptic compartments *in vivo* has yet to be determined. Existing RNA profiling approaches lack the spatial and temporal precision needed to resolve RNA repertoires restricted to cell-type specific synaptic compartments. To overcome this challenge, we developed PSD-95–Halo-seq, a light-induced proximity labelling technique that, when combined with long-read sequencing, selectively captures all post-synaptic full length RNA species in the excitatory post-synaptic compartment. Here, we applied this approach to investigate sex differences in RNA expression within excitatory synapses following exposure to an associative fear learning task in C57/Bl6 mice. We found dramatic sex differences in pseudogene and protein coding RNA expression, which are most abundant in females, with males exhibiting multiple noncoding RNA classes, including lncRNA, snoRNA, and rRNA. Females generally showed more 3′ UTR expression, and there was widespread differential exon usage following fear conditioning, including 179 isoforms in males, 69 in females, with no significant gene-level differential expression, indicating that behavioural state modifies sex-specific isoform usage rather than overall transcript abundance. Our discovery that synaptic RNA composition is dynamic, sexually dimorphic, and profoundly shaped by experience, offers new insight into previously inaccessible mechanisms underlying sex differences in fear-related learning and memory. PSD-95-Halo-seq is therefore a powerful method for the precise spatiotemporal identification of compartment-and cell-type–specific RNA.

**One sentence take-away:** Synapse-targeted proximity labelling and Long-Read sequencing demonstrate that excitatory post-synapses encode experience through sex-specific, isoform-level RNA remodeling.

## Introduction

Synaptic plasticity depends on the precise coordination of RNA-dependent regulatory processes operating across rapid timescales and highly restricted subcellular domains. Rather than serving as a passive intermediary, RNA actively shapes neuronal signaling through its roles in transcriptional output, alternative splicing, subcellular trafficking, localized translation, and RNA turnover. Neuron-specific splicing factors such as Nova, Rbfox, and PTBP proteins generate isoform repertoires that tune synaptic signaling (Ule et al., 2006; Raj & Blencowe, 2015). RNA localization is controlled by cis-elements and RNA-binding proteins, including ZBP1, FMRP, Staufen, and CPEB, which traffic transcripts into dendrites and maintain translational repression until synaptic activation (Doyle & Kiebler, 2011; Wang et al., 2016). Local translation is rapidly modulated by activity-dependent signaling pathways such as mTORC1, ERK–MAPK, and CaMKII (Kang & Schuman, 1996; Costa-Mattioli et al., 2009), while degradation pathways further refine transcript availability at individual synapses (Bolognani & Perrone-Bizzozero, 2004; Giorgi et al., 2007). Together, these processes position RNA as one of the critical substrates through which synapses encode activity and maintain plasticity.

The concept that synapses rely on local translation emerged from the discovery of polyribosomes beneath dendritic spines (Bodian, 1965; Steward & Levy, 1982) and was supported by evidence that disrupting dendritic translation impairs long-term potentiation and memory (Kang & Schuman, 1996; Sutton & Schuman, 2006). Activity-regulated transcripts, such as Arc, exemplify this principle: they undergo rapid nuclear induction, dendritic transport, and synapse-specific translation to modulate AMPA receptor trafficking and spine remodeling (Steward et al., 1998; Farris et al., 2014). Neurons maintain spatially segregated RNA repertoires whose abundance, isoform identity, and subcellular distribution shift dynamically with activity and experience (Holt & Schuman, 2013; Zhang et al., 2021). However, the full-length identities and isoform structures of RNAs residing directly within the post-synaptic density (PSD) remain unresolved in vivo.

Biological sex represents an additional, underexamined axis of synaptic molecular heterogeneity. Sex influences neuromodulator signaling, dendritic spine structure, chromatin dynamics, and activity-dependent gene regulation (Gillies & McArthur, 2010; Shansky, 2019). Large-scale transcriptomic analyses demonstrate pervasive sex differences in gene expression, RNA processing, and alternative splicing across cortical regions (Trabzuni et al., 2013; Khalaj et al., 2022), and recent work demonstrates sex-specific ribosome-associated RNA profiles in cortical neurons (Sare et al., 2021; Moran et al., 2022). Estrogen-dependent modulation of actin remodeling and glutamatergic signaling further suggests sex-dependent effects on post-transcriptional regulation within synapses (Oberlander & Woolley, 2016). Yet whether sex shapes the nanoscale RNA populations specifically embedded within excitatory—the molecular interface at which plasticity is expressed—remains unknown.

Here, we introduce PSD-95–Halo-seq, a proximity-labelling method that selectively tags RNAs within ∼100 nm of the PSD by fusing a HaloTag to a Fibronectin intrabodies generated with mRNA display (FingRs) against PSD-95. Combined with long-read sequencing, PSD-95–Halo-seq enables unbiased identification of full-length transcripts, isoforms, and RNA classes enriched at excitatory synapses in vitro and in vivo. Using this approach, we showed that the excitatory post-synapse harbors a distinct and diverse RNA repertoire as expected from mRNA local translation; that different sets of synaptic RNAs are found in animals of different sexes; and that behavioural experience drives isoform-level remodeling, that is also sex-dependent.

## Materials and methods

### Animals

Adult C57BL/6J mice (10–14 weeks; n = 6 males, n = 6 females) were used for all in vivo experiments. Animals were housed under a 12 h light/dark cycle with food and water ad libitum. All procedures followed the NHMRC Code for the Care and Use of Animals for Scientific Purposes and were approved by institutional ethics committees. Experimental design, stereotaxic targeting of prelimbic medial prefrontal cortex (mPFC), and behavioural handling followed previously published work from this laboratory (Marshall et al., 2024; Madugalle et al., 2023).

### Primary Neuronal Culture

Primary cortical neurons were prepared from E16–E17 embryos using a standard dissociation protocol adapted from Madugalle et al. (2023). Embryonic sex was determined using a combination of visual sexing and PCR-based genotyping of the Sry locus as described in McFarlane et al. (2013) (Fig.S1A). Neurons were plated on poly-D-lysine–coated dishes and maintained in Neurobasal medium supplemented with B27. For depolarization experiments, neurons were treated with 50 mM KCl for 30 mins over the 5mM present in media during standard culturing conditions. All experiments were conducted at 13/14 days in vitro (DIV 13/14).

### Lentiviral Constructs and Delivery

The PSD-95–HaloTag fusion construct was generated by inserting HaloTag7 into the C-terminus of FingR domain for PSD95 (Gross et al., 2013), with expression driven by the human Synapsin (hSyn) promoter. Membrane tethered mScarlet or mNeongreen is also expressed after a T2A sequence for identification of expressing cells. A Calm3 3’ UTR is inserted after the coding sequence before the WPRE sequence (Fig. S1B). Third-generation lentiviral particles were produced and concentrated using standard protocols for neuronal transduction, as described in Marshall et al. (2024).

For in vivo experiments, lentivirus expressing hSyn–PSD95–Halo was stereotaxically injected into the prelimbic mPFC (AP +1.9 mm, ML ±0.3 mm, DV −2.2 mm) using a Hamilton syringe at 50–75 nL/min. Injection volumes were 300–400 nL per hemisphere. Mice recovered for at least 2 weeks to allow stable expression before behavioural testing and Halo-seq labelling.

### Immunocytochemistry

Immunocytochemistry was performed based on Lo et al. (2022) on primary cortical neurons prepared as above on poly-D-Lysine-coated coverslips at low density and infected with PSD-95–HaloTag. At culture day 10 (DIV 10), cells were fixed in 4% formaldehyde prior to treatment with a fluorescent-conjugated Halo ligand (Janiela Fluor 646, Promega) and antibody treatment against PSD-95 (ab158258, Abcam at 1µg/mL). The coverslips were then mounted and imaged on a confocal microscope.

### Halo-seq Labelling In Vitro

Halo-seq labelling was performed based on Lo et al. (2022), with modifications to target synaptic compartments via PSD-95–Halo. Cultured neurons were incubated with the photosensitizer Dibromoflourescin (HaloDBF) for 15 minutes and subsequent incubation for 2 minutes with propargylamine prior to illumination with 530 nm light (Fig. S1C).

### In-situ visualization of labelled RNA

In-situ visualization was performed on primary cortical neurons based on Lo et al. (2022). Cultures were seeded as described for immunocytochemistry prior to labelling directed by PSD-95-HaloTag at DIV 10. Following labelling, an in-situ CuAAC click chemistry reaction was used to visualize labelled RNA with Alexa Fluor 568 dye (AF568 picolyl azide, Click Chemistry Tools 1292-1). Coverslips were then mounted and imaged on a confocal microscope.

### Halo-seq Labelling In Vivo

In vivo Halo-seq was adapted from the RNA proximity-labelling strategy described by Lo et al. (2022) for use in intact brain. 30 mins after fear conditioning or control handling, HaloDBF was infused bilaterally above the prelimbic mPFC through the existing craniotomy and incubated for 15 mins.

Propargylamine was similarly infused approximately 50 mins post fear conditioning or control handling. At 1-hour Animals were illuminated with 532 nm green via a fiberoptic placed in the existing craniotomy for 10 mins to oxidize RNA in the immediate vicinity of PSD-95–Halo–expressing synapses. Mice were then rapidly euthanized, and the mPFC was dissected for RNA extraction.

### RNA Extraction and Enrichment of Synaptic RNA

Total RNA was isolated using TRIzol or RNeasy kits according to the manufacturers’ instructions. Oxidized RNA was biotinylated by CuAAC click chemistry using a Dde Picoyl Azide–biotin reagent and enriched with streptavidin-coated magnetic beads, following the Halo-seq workflow of Lo et al. (2022) adapted for PSD-95–Halo targeting. Cleavage of the Dde linker with hydrazine facilitated release of RNA from the biotin-streptavidin complex suitable for Nanopore sequencing. Matched input (global) RNA was derived from the same cultures or animals prior to streptavidin bead treatment and processed for sequencing identically sans pulldown. RNA quality was assessed on an Agilent TapeStation; all samples used for sequencing had RIN > 8.

### Nanopore Library Preparation and Sequencing

Polyadenylated RNA was reverse-transcribed using the Oxford Nanopore cDNA-PCR kit. Sequencing libraries were prepared according to the manufacturer’s protocol and loaded onto R10.4 flow cells on MinION or GridION devices. Basecalling was performed using Guppy in super-accuracy mode.

Library preparation and depth were chosen to meet current best practices for long-read transcriptomics (Wang et al., 2021).

### Read Processing and Alignment

Reads with mean Q > 9 were aligned to the mouse genome (GRCm39) using minimap2 with splice-aware parameters optimized for long reads. Gene-level quantification was performed with featureCounts. Isoform reconstruction and quantification were performed using FLAIR or StringTie2 (Wang et al., 2021). RNA class and region annotations (e.g., protein-coding, lncRNA, pseudogene; exon, 5′ UTR, 3′ UTR) were assigned according to GENCODE vM29.

### Differential Expression Analyses Gene-level

DESeq2 was used to identify gene-level differential expression with Benjamini–Hochberg correction (*p*_*adj*_< 0.05). For in vitro experiments, the design formula was:

design = ∼ sex + treatment + fraction (synapse vs input) For in vivo experiments, the design formula was:

design = ∼ sex + conditioning + fraction

#### Isoform-level

Differential exon usage and isoform regulation were assessed using DEXSeq. Isoforms with *p*_*adj*_< 0.05 and supported by ≥20 reads across samples were considered significant.

#### Gene Ontology and RNA Class Analyses

Gene Ontology enrichment was performed using clusterProfiler. RNA class and region analyses were based on GENCODE vM29 annotations, with counts aggregated by RNA biotype and by genomic region (exon, 5′ UTR, 3′ UTR, intron).

#### Fear Conditioning

Fear conditioning was carried out as in prior work from this laboratory (Madugalle et al., 2023; Marshall et al., 2024). Briefly, mice were habituated and then exposed to 5 presentations of a 120s light cue co-terminating with a 0.7 mA, 1s foot shock. Freezing behaviour was recorded using FreezeFrame and analyzed offline. Halo-seq labelling was initiated 1 h after training to target the early consolidation phase of memory formation.

### Statistical Analysis

All statistical analyses were conducted in R, Python or GraphPad Prism. Data normality was assessed with Shapiro–Wilk tests and homogeneity of variance with Levene’s or Bartlett’s tests. Multi-group comparisons were performed using two-way or three-way ANOVA as appropriate, followed by Benjamini–Hochberg–corrected post hoc comparisons. Exact n values, statistical tests, F statistics, and p-values are reported in the Results and figure legends. Experimenters were blind to sex and behavioural condition during tissue collection, RNA processing, and sequencing.

### Data and reagent availability

All data contained within figures is available in supplemental datasets. All custom scripts used are available at https://github.com/GenomicPlasticityLab/HaloSeq. All constructs are available from Addgene at https://www.addgene.org/plasmids/articles/28263779/250667/. All commercial reagents are listed and any custom reagents available on request.

## Results

### PSD-95–Halo-seq enables targeted labelling of post-synaptic RNA in neurons

To achieve nanometer-scale resolution of synaptic RNA, we adapted Halo-seq for post-synaptic targeting by fusing a HaloTag and FingR domain to PSD-95 (Gross et al., 2013) and expressing it under the human Synapsin promoter (hSyn). A Calm3 3’ UTR, was also included to assist in trafficking to the dendrites. This configuration in entirety confines light-inducible oxidative RNA labelling to excitatory post-synapses (Fig. 1A). In cultured mouse cortical neurons **(n = 3 cultures)**, viral expression of PSD-95–Halo produced punctate HaloTag signal in dendrites and spines that colocalized with endogenous PSD-95, confirming accurate synaptic localization (Fig. 1B). However, we still observe signal throughout the soma which may represent somatic synapses or nascent PSD-95 yet to be trafficked to the dendrites.

**Figure 1:**
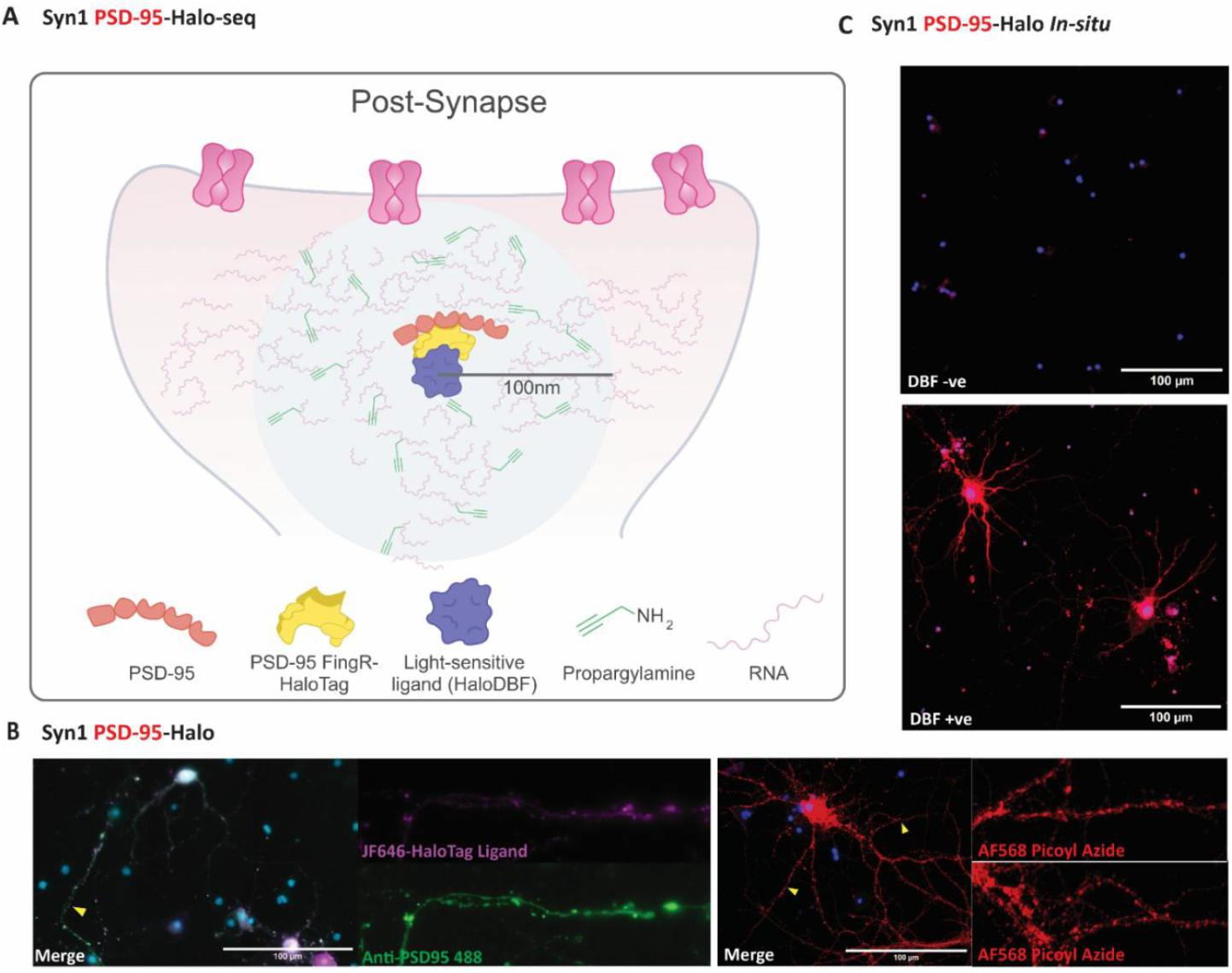
Halo-seq facilitates the labelling of post-synaptic RNA populations. **A.** Schematic outlines the Syn1-PSD95-Halo-seq labelling methodology.**B.** Immunocytochemistry validation of halo-tag colocalization with PSD95 protein in primary cultured neurons. HaloTag localization visualized with fluorescent dye-conjugated halo ligand (JF646 HaloTag Ligand). **C.** in-situ labelling of proximal RNA visualized though CuAAC click chemistry of a fluorescent dye in the presence and absence of HaloDBF. (DBF-, n = 3 and DBF+, n = 3). Yellow icons denote the location of magnified panels

To visualize RNA labelling in situ, neurons were incubated with the photosensitizer HaloDBF, illuminated with green light, and labelled RNA was conjugated to a fluorescent azide via CuAAC click chemistry. This generated discrete dendritic puncta consistent with post-synaptic RNA populations, whereas labelling was absent in the absence of HaloDBF (Fig. 1C). These observations confirm that PSD-95–Halo targets nanometer-spaced RNA labelling specifically to post-synaptic compartments.

Biotinylation of alkylated RNA followed by streptavidin blotting showed strong signal exclusively when all components—HaloTag, HaloDBF, light, and propargylamine—were present following optimization with cleavable Dde Biotin Picoyl Azide (Figure S2), demonstrating specificity of the oxidative labelling reaction.

### PSD-95–Halo-seq identifies a synapse-enriched transcriptome in vitro

To define the composition of post-synaptic RNA, we performed PSD-95–Halo-seq on cultured cortical neurons under four conditions: male or female neurons, each exposed either to basal activity or to depolarization with 50 mM KCl (Fig. 2A; **n = 6 biological replicates per group**). Synapse-labelled RNA was isolated through streptavidin pulldown and compared with paired global RNA inputs. All samples were poly(A)-tailed and sequenced using Oxford Nanopore long-read sequencing to resolve full-length transcripts and isoforms.

**Figure 2:**
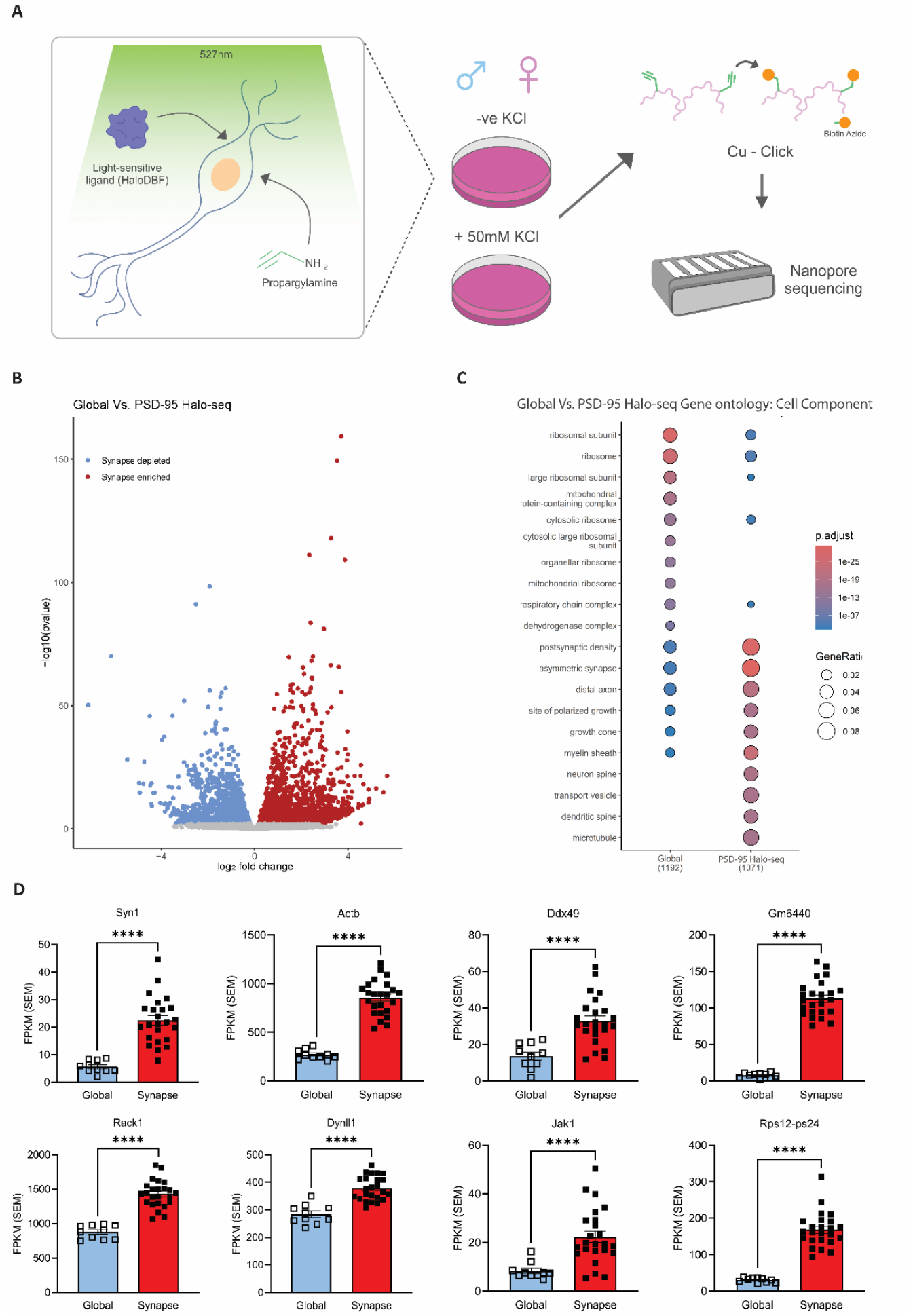
Identifying post-synaptic transcriptomes with PSD-95 Halo-seq. **A.** Schematic detailing the experimental pipeline followed to derive post-synaptic RNA sequence data. **B.** Volcano plot showing differentially expressed genes comparing global RNA against PSD-95 halo-seq capture at the post-synapse. Significance is determined by DeSeq2 with an adjusted *p-*value threshold of ≤0.05. Significant, synapse enriched genes are shown in red. **C.** Gene ontology analysis of global and synapse enriched genes where input gene lists were restricted to differentially expressed genes output from DeSeq2 with *p*_*adj*_< 0.05 and a corresponding geneID within the genome annotation file used. Cell component terms are given as the top 10 most significant for each group. **D.** Bar plots showing representative transcripts (Syn1 t (29) = 9.026, p = 6.3 × 10^-10^; Actb t (30) = 9.026, p = 3.0 × 10^-15^; Ddx49 t (30) = 5.574, p = 4.52 × 10^-6^; Gm6440 t (25) = 21.06, p = 1.0 × 10^-15^; Rack1 t (31) = 10.93, p = 4.05 × 10^-12^; Dynll1 t (30) = 5.574, p = 1.73 × 10^-6^; Jak1 t (30) = 5.493, p = 5.6 × 10^-6^; Rps12-ps24 t (25) = 13.95, p = 2.82 × 10^-13^) that are significantly enriched at the post-synapse expressed as Fragments Per Kilobase of transcript per Million mapped reads (FPKM). For every gene comparison there was n = 10 for Global and n = 24 for Synapse. Data were analyzed by unpaired Welch’s t-test: \*\**p* < 0.01, \*\*\**p* < 0.001, ****p <0.0001. bars show Mean +/-SEM, individual cultures shown as discrete points.)

Pooled synapse-enriched RNA was contrasted with global RNA using DESeq2 (negative binomial model; *p*_*adj*_< 0.05, Benjamini–Hochberg). Out of 28,433 detected genes, 3,299 showed significant differential expression, with 1,927 enriched and 1,372 depleted at the synapse (Fig. 2B).

Gene Ontology (GO) analysis of enriched transcripts revealed overrepresentation of synaptic and perisynaptic compartments, including the post-synaptic density (FDR < 10^−4^), glutamatergic synapse, axon terminus, and periactive zone (Fig. 2C). These findings validate that PSD-95–Halo-seq isolates molecularly authentic post-synaptic RNA populations.

Synapse-enriched RNA was also compared with DESeq2 by sex and KCl treatment, however few genes displayed differential expression (Table S1)

Several canonical synaptic transcripts were highly enriched, including *Syn1* (t (29) = 9.026, p = 6.3 × 10^-10)^ and *Rack1* (t (31) = 10.93, p = 4.05 × 10^-12^), scaffolding proteins essential for synaptic structure and signaling (Fig. 2D). Cytoskeletal regulators such as *Actb* (t (30) = 9.026, p = 3.0 × 10^-15^) and the dynein light chain *Dynll1* (t (30) = 5.574, p = 1.73 × 10^-6^) were also enriched, consistent with local cytoskeletal remodeling at active synapses. Additional enriched transcripts included *Jak1* (t (30) = 5.493, p = 5.6 × 10^-6^), a kinase involved in JAK/STAT signaling and synaptic plasticity.

Long-read sequencing revealed abundant ribosomal-protein pseudogene transcripts in the synaptic fraction, including *Rps* and *Rpl* pseudogene families. Although traditionally classified as nonfunctional, pseudogenes can regulate translation through miRNA sequestration or competitive binding with cognate transcripts. Their high abundance here suggests a potential regulatory role for pseudogenes in synaptic translation.

### Sex and activity state determine RNA class and isoform abundance at the post-synapse in vitro

To quantify the influence of sex and KCl-induced depolarization on synaptic RNA composition, we annotated significantly synapse-enriched transcripts by RNA class (coding, lncRNA, pseudogene, snoRNA, etc.). A two-way ANOVA revealed significant effects of RNA class (F (7, 160) = 921.4, 1.0 × 10^-15^), and class × group interaction (F (21, 160) = 4.35, p = 3.13 × 10^-8^) (Fig. 3A). Post-hoc tests (BH-corrected) showed major differences in protein-coding (Male KCl q(160) = 8.544, p = 6.19 × 10^-8^, Female KCl q(160) = 3.797, p = 0.0397, KCl-q(160) = 5.221, p = 0.0017, KCl+ q(160) = 7.12, p = 7.58 × 10^-6^) and pseudogene (Male KCl q(160) = 8.781, p = 2.64 × 10^-8^, Female KCl q(160) = 3.797, p = 0.0396, KCl-q(160) = 5.458, p = 0.0009, KCl+ q(160) = 7.12, p = 7.58 × 10^-6^) abundance between male vs female neurons and between KCl-treated vs untreated cultures.

**Figure 3:**
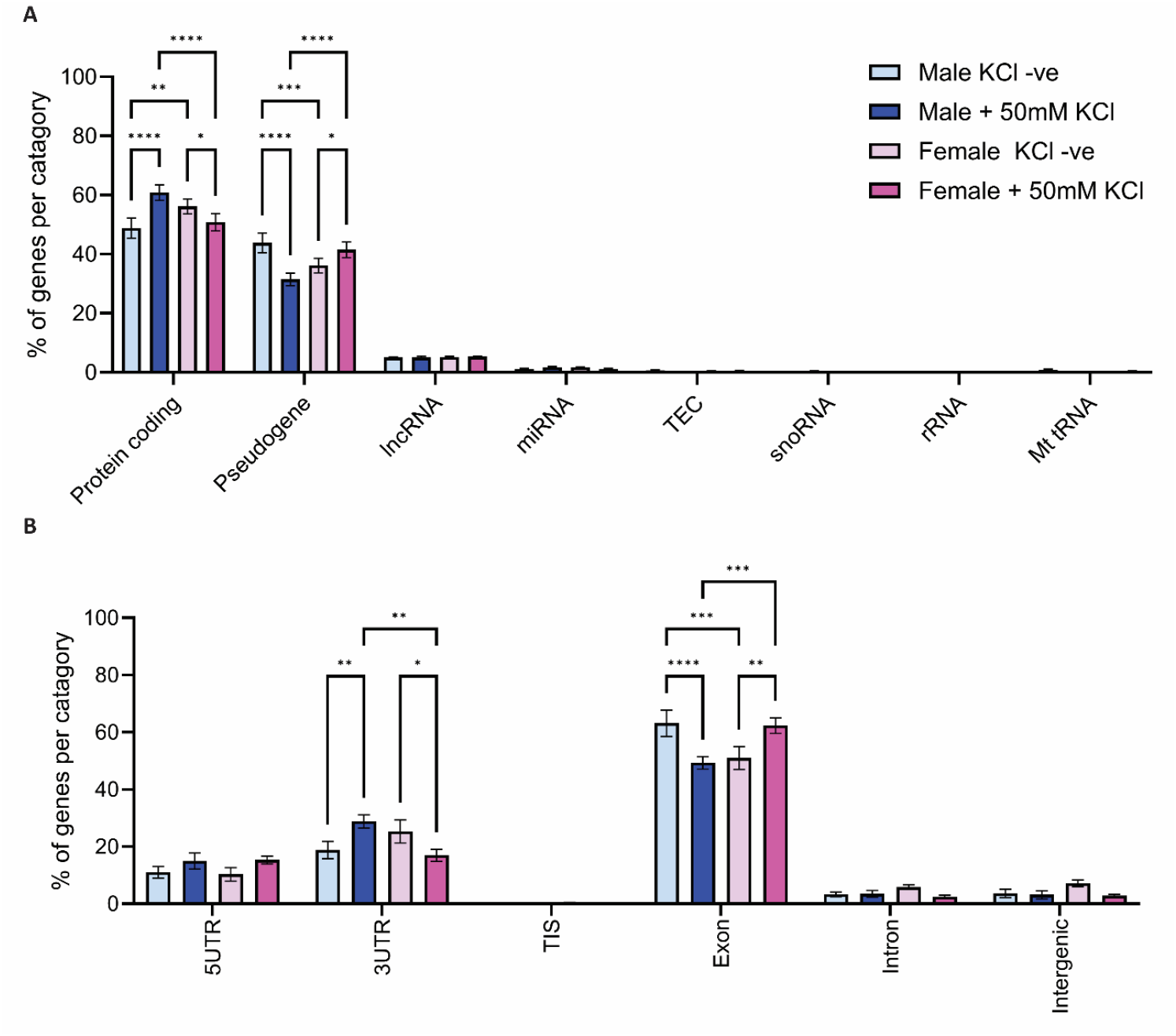
Post-synaptic transcriptomes are dynamic according to sex and activations state. **A.** Bar plots showing % total abundance of synapse-enriched transcripts across sex and treatment with 50 mM KCl by RNA class (Class F(7, 160) = 921.4, p = 1.0 × 10^-15^; class × group interaction (F(21, 160) = 4.35, p = 3.13 × 10^-8^) Post-hoc tests: protein-coding, Male KCl q(160) = 8.544, p = 6.19 × 10^-8^, Female KCl q(160) = 3.797, p=0.0397, KCl-q(160) = 5.221, p = 0.0017, KCl+ q(160) = 7.12, p = 7.58 × 10^-6^) and pseudogene (Male KCl q(160) = 8.781, p = 2.64 × 10^-8^, Female KCl q(160) = 3.797, p = 0.0396, KCl-q(160) = 5.458, p = 0.0009, KCl+ q(160) = 7.12, p = 7.58 × 10^-6^). **B.** Bar plots showing % total abundance of synapse-enriched transcripts across sex and KCl treatment by transcript component (Component F(5, 120) = 375.3, p = 1.0 × 10^-15^; class × group interaction (F(15, 120) = 4.084, p = 5.54 × 10^-6^) Post-hoc tests: 3UTR, Male KCl q(120) = 4.591, p = 0.0082, Female KCl q(120) = 3.826, p = 0.0386, KCl+ q(120) = 5.432, p = 0.0011 and exon (Male KCl q(120)=6.351, p = 9.6 × 10^-5^, Female KCl q(120) = 5.203, p = 0.002, KCl-q(120) = 5.585, p = 0.0008, KCl+ q(120) = 5.968, p = 0.0003) (Group, n = 6, bars show Mean +/-SEM) (* *p* < 0.05,** *p* < 0.01,*** *p* < 0.001,**** *p* < 0.0001).

**Table 1:** Neuronal activation generates shifts in isoform abundance at the post-synapse. Genes identified from PSD-95-Halo-seq samples with DEXseq showing differential exon usage between groups. Significant Isoforms demonstrated *p*_*adj*_< 0.05 and were supported by ≥20 reads across samples.

| Comparison | Gene ID | Gene | Description | Isoform Function |
| --- | --- | --- | --- | --- |
| Female Vs. Female + 50mM KCl treatment | ENSMUSG00000071369 | *Map3k5* | Major component of the MAPK signaling cascade. Map3k5 receives signals from upstream proteins and initiates the pathway. | Isoforms influence downstream signaling specificity (Maik-Rachline, Wortzel, Seger 2021). |
| Female Vs. Female + 50mM KCl treatment | ENSMUSG00000060402 | *Chst8* | Encodes the N-acetylglucosamine-6-sulfotransferase 8 (GlcNAc-6ST) enzyme, which sulfates glycans on proteins | Multiple coding and non-coding isoforms, distinct function unknown. |
| Female Vs. Female + 50mM KCl treatment | ENSMUSG00000035232 | *Pdk3* | A mutation in the PDK3 gene has been linked to impaired synaptic transmission, as this mutation affects ATP production. | Three isoforms, function of non-coding isoforms unknown. |
| Female Vs. Female + 50mM KCl treatment | ENSMUSG00000026880 | *Stomatin* | Encodes a membrane protein which interacts with the postsynaptic side of synapses by forming complexes that regulate ion channels, especially acid-sensing ion channels (ASICs). | Two main isoforms, one containing a retained intron, speculatively for regulation of translation. |
| Female Vs. Female + 50mM KCl treatment | ENSMUSG00000020176 | *Grb10* | Encodes an adapter protein interacting with growth factor receptors for insulin and leptin for example. Found in post synapse to regulate synaptic function. | High level of splicing, Splice variants of human gene implicated in mitogen signaling / insulin growth factor 1, mouse unknown (O'Neill et al., 1996). |
| Male Vs. Male + 50mM KCl treatment | ENSMUSG00000025278 | *Flnb* | Encodes actin-binding protein important for linking cell membrane to cytoskeleton, including at synapse. It acts as a scaffold for *Rac1* signaling and has been shown to regulate alternative splicing in certain contexts. | Two main isoforms shown to have tissue-specific abundance. Isoform differential function unknown. |

Overrepresentation analysis of RNA regions also revealed significant group-dependent shifts in RNA transcript component (Component F (5, 120) = 375.3, p = 1.0 × 10^-15^; class × group interaction (F (15, 120) = 4.084, p = 5.54 × 10^-6^; Fig.3B). Specifically, Females displayed significantly less 3′ UTR following KCl stimulation (q (120) = 3.826, p = 0.0386), whereas males displayed significantly more (q (120) = 4.591, p = 0.0082). Exon usage mirrored this with females displaying significantly more exon after stimulation (q (120) = 5.203, p = 0.002), and males significantly less (q (120) = 6.351, p = 9.6 × 10^-5^). These results indicate that **sex explains more variance in synaptic RNA architecture than activation state**.

To assess isoform-level regulation, we performed DEXSeq on synapse-enriched RNA. Five RNAs showed significant differential exon usage (*p*_*adj*_< 0.05) in females following KCl stimulation, including *Map3k5, Grb10, Chst8, Stomatin*, and *Pdk3*. These genes participate in MAPK signaling, ion channel regulation, and membrane trafficking—pathways known to modify synaptic responsiveness.

In contrast, only a single gene (*Flnb*, Filamin B) exhibited differential exon usage in males. Notably, none of these isoform changes were accompanied by significant gene-level differential expression in DESeq2, indicating that activity primarily regulates the structure, not the abundance, of synaptic transcripts.

Together, these results demonstrate that sex and activity state exert strong, isoform-specific effects on post-synaptic RNA architecture.

### PSD-95–Halo-seq profiles synaptic RNA in vivo during memory formation

We next applied PSD-95–Halo-seq in the medial prefrontal cortex (mPFC) of freely behaving adult mice (**n = 6 males, n = 6 females**). Animals underwent cued fear conditioning, and synaptic RNA labelling occurred 1 h after training—a critical window for memory consolidation (Fig. 4A).

**Figure 4:**
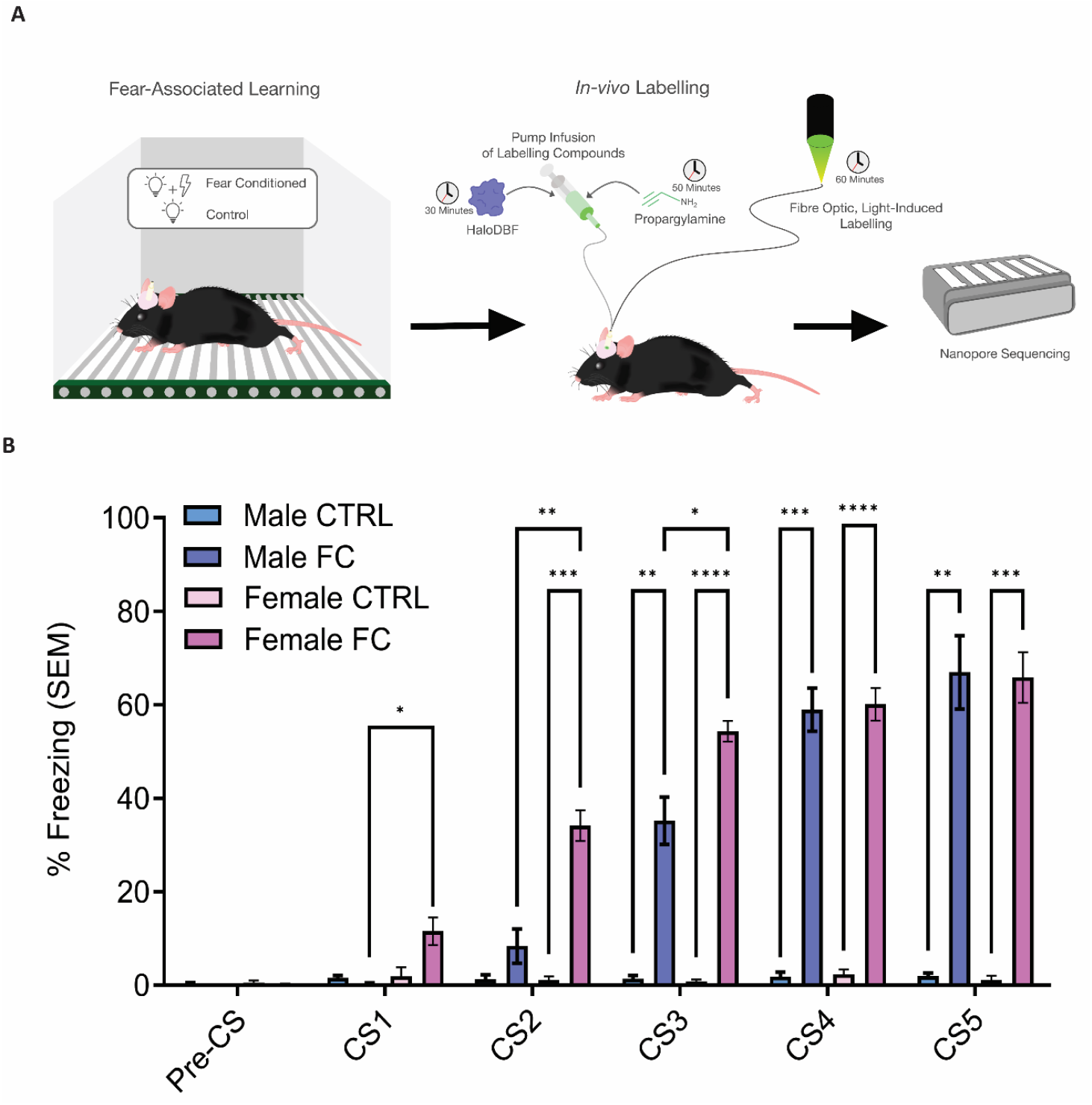
Identifying post-synaptic transcriptomes in-vivo with PSD-95 Halo-seq. **A.** Schematic detailing the experimental pipeline, including timeline for labelling in-vivo given as minutes post fear learning. **B** Fear training resulting in a significant increase in freezing over time (n = 6/group, three-way ANOVA CS F(2, 49) = 116.4, p =1 × 10^-15^), Post-hoc tests: CS1 (FC q(5.041) = 5.4, p = 0.0435), CS2 (q(9.881) = 7.4, p = 0.0019, Female q(5.62) = 13.82, p = 0.0004), CS3 (FC q(6.817) = 4.895, p = 0.0426, Female q(5.315) = 33.89, p = 6.61 × 10^-6^, Male q(5.216) = 9.349, p = 0.0039) CS4 (Female q(5.84) = 22.63, p = 1.9 × 10^-5^, Male q(5.456) = 17.09, p = 0.0002) and CS5 (q(5.244) = 16.75, p = 0.0002, Male q(5.065) = 11.68, p = 0.0015). Mean +/-SEM are shown in B. * *p* < 0.05, ** *p* < 0.01, *** *p* < 0.001, **** *p* < 0.0001.

Behaviourally, females appeared to acquire the task more rapidly as shown by significantly higher freezing at CS2 (q (7) = 9.881, p = 0.0019) and CS3 (q (7) = 4.895, p = 0.0426), though both conditioned groups had the significant changes to their controls, and non-significant freezing differences by the last CS (Fig. 4B). Synaptic samples were then enriched from the mPFC of these animals and compared with paired global RNA from each animal.

### Global vs. synaptic enrichment in vivo

Long-read sequencing identified 27,939 transcripts in total, with 2,815 significantly different between synaptic and global fractions. Of these, 1,684 were synapse-enriched and 1,132 depleted (*p*_*adj*_< 0.05; Fig. 4B). GO enrichment again highlighted post-synaptic density, asymmetric synapse, and glutamatergic synapse as top-ranked terms (FDR < 10^−4^), verifying synaptic specificity in vivo (Fig. 4C).

Representative enriched transcripts included *Vamp2* (t (15) = 6.495, p = 9.19 × 10^-6^), *Grin2a* (t (19) = 4.256, p = 0.0004), *Slc1a2* (t (14) = 7.214, p = 4.01 × 10^-6^), *Nsmf* (t (13) = 7.285, p = 5.09 × 10^-6^), *Sptbn2* (t (13) = 7.714, p = 3.71 × 10^-6^), and *Dynll2* (t (22) = 2.741, p = 0.0120) (Fig. 4D), reflecting synaptic vesicle cycling, ionotropic glutamate receptor signaling, and cytoskeletal regulation. In contrast to the in vitro dataset, pseudogene enrichment was markedly reduced in vivo.

Synapse-enriched RNA was also compared by sex and conditioning with DESeq2, revealing sex to be another determinate of transcript abundance (Table S2).

### Sex is the dominant determinant of RNA class composition in vivo

Annotating synapse-enriched transcripts by RNA class revealed pronounced sex differences (Fig. 5A). A two-way ANOVA showed significant effects of RNA class (F (13, 112) = 191.6, p = 1.0 × 10^-15^) and interaction (F (39, 112) = 26.61, p = 1.0 × 10^-15^). Post-hoc comparisons (BH-corrected) indicated that females uniquely displayed high pseudogene (Control q (112) = 15.56, p = 6.2 × 10^-14^, Conditioned q (112) = 22.82, p = 4 × 10^-14^) and protein coding (Control q (112) = 15.79, p = 5.9 × 10^-14^, Conditioned q (112) = 12.4, p = 2.15 × 10^-13^) abundance, whereas males were enriched for multiple noncoding RNA classes including lncRNA (Control q (112) = 12.3, p = 2.75 × 10^-13^, Conditioned q (112) = 12.99, p = 9.1 × 10^-14^), snoRNA snoRNA (Control q (112) = 15.44, p = 6.3 × 10^-14^, Conditioned q (112) = 15.06, p = 6.9 × 10^-14^), and rRNA (Control q (112) = 4.894, p = 0.0042, Conditioned q (112) = 5.839, p = 0.0004).

**Figure 5:**
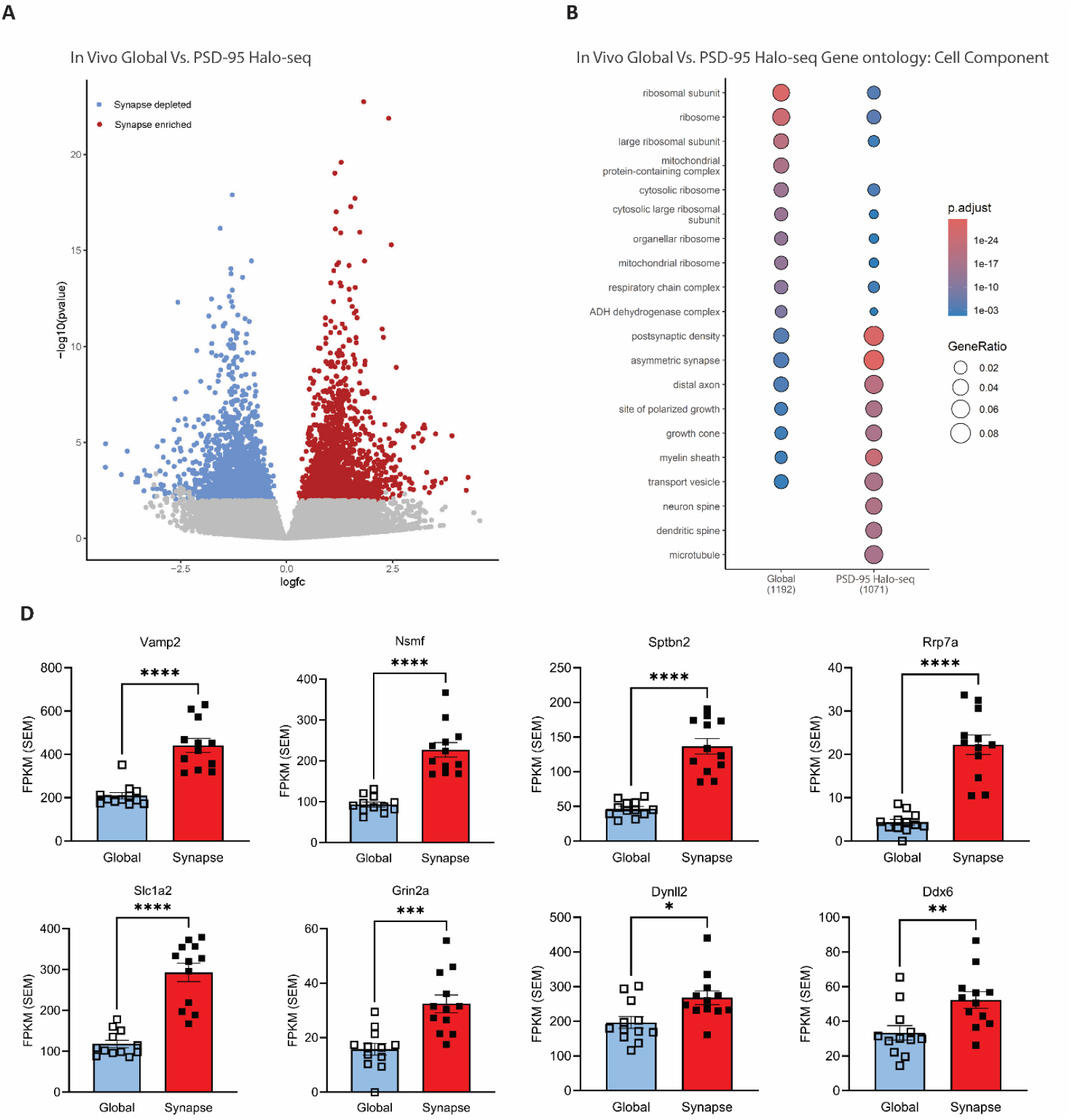
Identifying post-synaptic transcriptomes in-vivo with PSD-95 Halo-seq. **A.** Volcano plot showing differentially expressed genes comparing global RNA against PSD-95 halo-seq capture at the post-synapse. Significance is determined by DeSeq2 with an adjusted *p-*value threshold of ≤0.05. Significant, synapse enriched genes are shown in red. **B.** Gene ontology analysis of global and synapse enriched genes where input gene lists were restricted to differentially expressed genes output from DeSeq2 with *p*_*adj*_< 0.05 and a corresponding geneID within the genome annotation file used. Cell component terms are given as the top 10 most significant for each group. **C.** Bar plots showing representative transcripts (Vamp2 t(15) = 6.495, p = 9.19 x 10^-6^; Nsmf t(13) = 7.285, p = 5.09 × 10^-6^; Sptbn2 t(13) = 7.714, p = 3.71 × 10^-6^; Rrp7a t(13) = 7.695, p = 3.55 × 10^-6^; Slc1a2 t(14) = 7.214, p = 4.01 × 10^-6^; Grin2a t(19) = 4.256, p = 0.0004; Dynll2 t(22) = 2.741, p = 0.0120; Ddx6 t(22) = 3.002, p = 0.0066) that are significantly enriched at the post-synapse expressed as Fragments Per Kilobase of transcript per Million mapped reads (FPKM). For every gene comparison there was n = 12 for Global and n = 12 for Synapse. Data were analyzed by unpaired Welch’s t-test: **p < 0.01, ***p < 0.001, ****p <0.0001. Bars show Mean +/-SEM, individual animals shown as discrete points.

Overrepresentation analysis of RNA regions also revealed significant group-dependent shifts in RNA transcript component (Component F (5, 48) = 388.5, p = 1.0 × 10^-15^; class × group interaction F (15, 48) = 12.35, p = 1.01 × 10^-11^; Fig.5B). Specifically, females generally displayed significantly more 3’UTR (Control q (48) = 9.408, p = 1.49 × 10^-7^, Conditioned q (48) = 7.557, p = 1.45 × 10^-5^) and less exon (Control q (48) = 12.18, p = 1.54 × 10^-10^, Conditioned q (48) = 6.015, p = 0.0005) usage compared to males. Additionally, fear conditioning induced an increase in exon usage, only in females (Female q (48) = 5.706, p = 0.0011). These results indicate that **sex explains more variance in synaptic RNA architecture than behavioural state**.

### Learning drives sex-specific isoform remodeling in vivo

DEXSeq identified widespread differential exon usage following fear conditioning (**179 isoforms in males, 69 in females**, *p*_*adj*_< 0.05; Table 2). Surprisingly, DESeq2 detected **no significant gene-level differential expression**, indicating that behavioural state modifies isoform usage rather than overall transcript abundance.

Key genes exhibiting isoform changes included *Dlg2, Camk2b* and *Fos* whose splice variants influence synaptic adhesion, ion channel localization, and membrane scaffolding. These findings suggest that isoform remodeling is a major regulatory axis during memory consolidation and that males and females engage distinct molecular programs at the PSD.

## Discussion

This study establishes PSD-95–Halo-seq as a high-resolution approach for defining the RNA architecture of excitatory synapses. By combining nanometer-scale proximity labelling with long-read sequencing, we resolve full-length transcripts, isoforms, and RNA classes directly within the post-synaptic density (PSD)—a subcellular compartment that has remained largely inaccessible at isoform-level resolution *in vivo*. Significantly, this approach improves the resolution of synaptic RNA capture over existing methods (Davies, 2025 NLM) and compliments the use of PSD-95 proximity-labelling for local proteome capture (Uezu et al., 2016).

The enrichment of canonical synaptic and dendritic transcripts validates the specificity of PSD-95– Halo-seq and aligns with the long-standing observation that local translation underlies enduring forms of synaptic plasticity (Steward & Levy, 1982; Kang & Schuman, 1996; Sutton & Schuman, 2006). We chose to use the FingR domain targeting PSD95 instead using full-length PSD95 or truncated PSD95 to avoid any possible adverse effects associated with PSD95 overexpression that had been reported previously (Zhang et al., 2012; Dore et al., 2021).

A central finding is that **sex is a dominant determinant of synaptic RNA composition and isoform structure**. While sex differences in transcription, RNA processing, and translation have been documented in cortical tissue and ribosome-associated populations (Trabzuni et al., 2013; Sare et al., 2021; Moran et al., 2022), our results extend these effects to excitatory synapses. In vivo, females showed markedly greater enrichment of pseudogenes, whereas males exhibited higher representation of several noncoding RNA classes, indicating that sex biases are not uniform but transcript-type specific. This suggests that sex shapes post-transcriptional regulation at synapses through mechanisms that may include miRNA competition, differential exon usage, and RNA binding protein (RBP)-mediated control of translation, although these pathways remain to be directly tested in this compartment. Together, the findings demonstrate that the molecular substrates supporting synaptic plasticity and memory differ substantially between male and female, positioning sex as a major source of synaptic nanoscale transcriptomic diversity.

Experience **further remodels the excitatory synaptic transcriptome in a sex-dependent manner**. Fear conditioning induced robust isoform-specific changes in both males and females, yet these occurred largely in the absence of gene-level differential expression. This dissociation indicates that isoform structure—rather than transcript abundance—is the primary dynamic layer of post-synaptic regulation during consolidation. Several of the behaviourally regulated isoforms we identified, including variants of *Camk2b, Dlg2, Fos, Map3k5, Grb10*, and *Stomatin*, map onto pathways linked to cytoskeletal remodeling, excitatory receptor signaling, and synaptic adhesion, suggesting that experience selectively tunes synaptic function through transcript architecture. These findings are consistent with emerging models in which plasticity relies on rapid, isoform-level reconfiguration of neuronal transcripts rather than broad transcriptional activation, enabling synapses to modulate protein localization, interactions, and signaling specificity with high temporal and spatial precision.

The prominence of isoform-level regulation raises an additional mechanistic question: whether these isoforms arise from local processing within the synapse itself. One interpretation of the predominance of isoform-specific changes is the possibility of local splicing within dendrites and synapses. Although our data do not directly demonstrate subcellular splicing, multiple studies have identified spliceosomal components, pre-mRNA processing machinery, and alternatively spliced transcripts in dendritic compartments, beginning with the initial report of synaptic spliceosome markers (Glanzer et al., 2005) and supported by subsequent evidence of dendritic U2 snRNP localization, localized exon inclusion, and synaptic splice factor regulation (Bell et al., 2008; Bell et al., 2010; Ortiz et al., 2017; Thomas-Jinu et al., 2017). The isoform patterns we observe—particularly those involving signaling adaptors and ion-channel modulators—are consistent with such mechanisms. Additionally, preliminary proteomics from synaptosomes of trained animals (not shown) indicate the presence of spliceosomal proteins in the mPFC, further supporting synaptic-level RNA processing as a plausible contributor. Future studies that causally manipulate splicing within defined synaptic compartments will be necessary to test this lingering but compelling hypothesis.

A comparison of the in vitro (Fig. 3) and in vivo (Fig. 6) datasets highlights how neuronal context shapes post-synaptic RNA composition. The strong pseudogene enrichment observed in cultured neurons was markedly attenuated in vivo, suggesting that dissociated cultures amplify certain RNA regulatory pathways or that intact circuits impose tighter control over noncoding RNA abundance. These differences are consistent with prior demonstrations that neuronal culture alters RNA transport dynamics, RBP availability, and translation-associated regulatory programs relative to the intact brain (Wang et al., 2016; Holt & Schuman, 2013). More broadly, the divergence between cultured and in vivo synaptic transcriptomes underscores the importance of studying RNA regulation within behaviourally engaged neural circuits, where hormonal state, neuromodulatory tone, and activity patterns more accurately reflect the physiological pressures acting on synaptic RNA populations.

**Figure 6:**
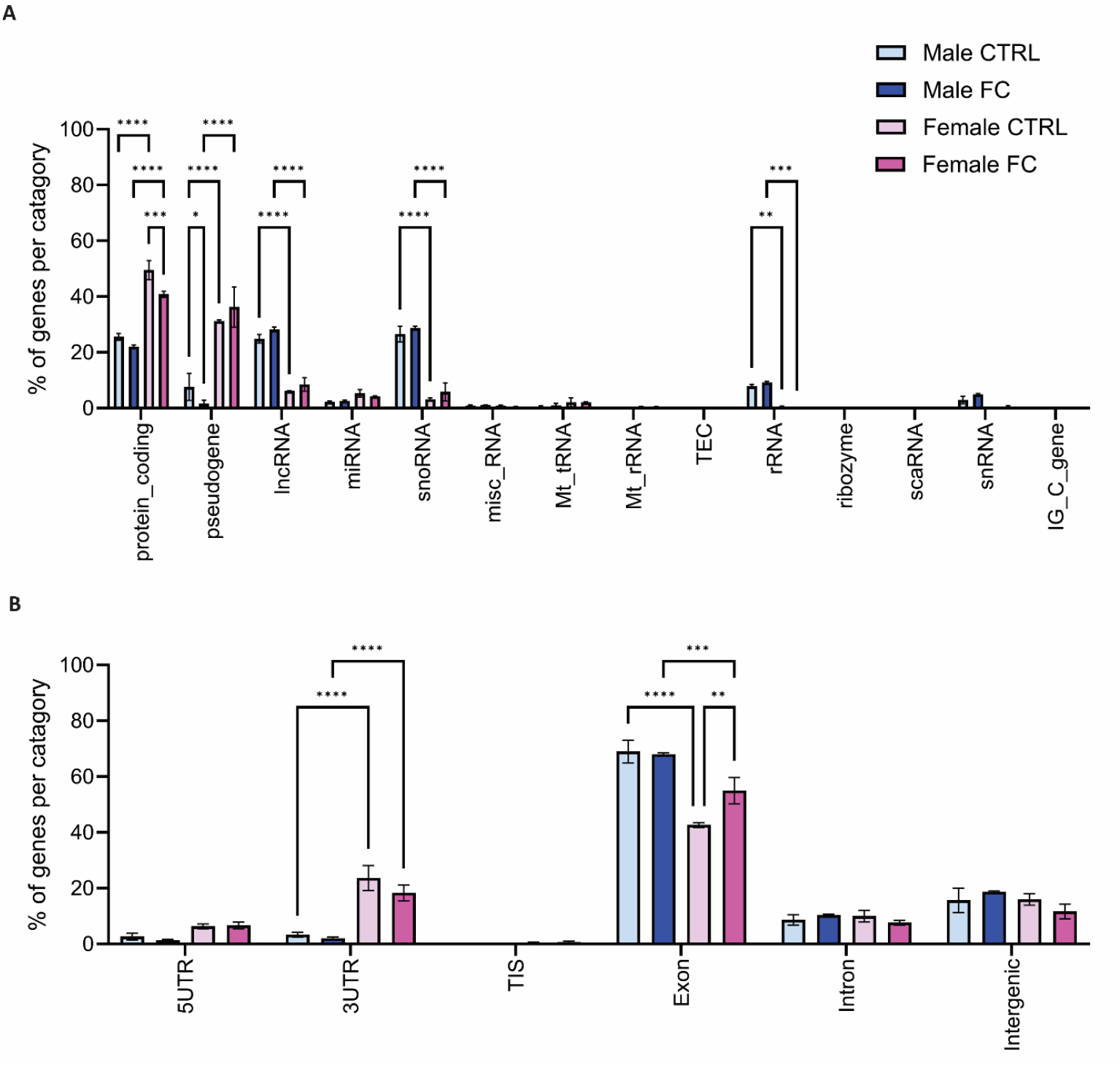
In-vivo post-synaptic transcriptomes dynamically encode experience in a sex-specific manner. **A.** Bar plots showing % total abundance of synapse-enriched transcripts across sex and experience by RNA class (Class F(13, 112) = 191.6, p = 1.0 × 10^-15^; class × group interaction F(39, 112) = 26.61, p = 1.0 × 10^-15^). Post-hoc tests: protein-coding, Female q (112) = 5.736, p = 0.0005, Control q (112) = 15.79, p = 5.9 × 10^-14^, Conditioned q (112) = 12.4, p = 2.15 × 10^-13^, pseudogene, Male q (112) = 3.942, p = 0.0313, Control q (112) = 15.56, p = 6.2 × 10^-14^, Conditioned q (112) = 22.82, p = 4 × 10^-14^, lncRNA, Control q (112) = 12.3, p = 2.75 × 10^-13^, Conditioned q (112) = 12.99, p = 9.1 × 10^-14^, snoRNA, Control q (112) = 15.44, p = 6.3 × 10^-14^, Conditioned q (112) = 15.06, p = 6.9 × 10^-14^, and rRNA, Control q (112) = 4.894, p = 0.0042, Conditioned q (112) = 5.839, p = 0.0004. **B.** Bar plots showing % total abundance of synapse-enriched transcripts across sex and KCl treatment by transcript component (Component F (5, 48) = 388.5, p = 1.0 × 10^-15^; class × group interaction F (15, 48) = 12.35, p = 1.01 × 10^-11^). Post-hoc tests: 3UTR, Control q (48) = 9.408, p = 1.49 × 10^-7^, Conditioned q (48) = 7.557, p = 1.45 × 10^-5^ and exon, Female q (48) = 5.706, p = 0.0011, Control q (48) = 12.18, p = 1.54× 10^-10^, Conditioned q (48) = 6.015, p = 0.0005 (Group, n = 3, Error bars represent SEM) (* *p* < 0.05,** *p* < 0.01,*** *p* < 0.001,**** *p* < 0.0001).

### Limitations

PSD-95–Halo-seq relies on oxidative tagging, which may preferentially modify exposed or single-stranded regions of RNA, potentially introducing structural biases in capture efficiency. Furthermore, because Halo-seq depends on oxygen-driven photochemistry, factors such as local illumination, tissue scattering, and oxygen availability may produce variability in labelling efficiency in vivo. To address these constraints, *in vivo–*compatible organic electronic illumination platforms are being actively developed in collaboration with co-author Assoc.Prof. Marcin Kielar, whose recent work demonstrates the feasibility of organic light-emitting devices (OLEDs) for neuronal photonic stimulation (Kielar et al., 2022). In this context, implantable OLED-based deep-tissue stimulators can bring the advantages of color-tunability, ultra-small footprint, flexibility and biocompatibility, representing significant improvements on current optical techniques in the field (Kielar et al., 2025). Additionally, PSD-95 expression is predominant but not exclusive to the post-synapse, necessitating the use of a global RNA control to contrast populations; potentially reducing detection of post-synaptic RNA not enriched. Finally, although long-read sequencing enables isoform-level resolution, sequencing depth remains a limiting factor for detecting rare or transient transcript isoforms.

### Future Directions

The modularity of Halo-seq permits flexible targeting of different synaptic proteins, enabling systematic comparison of the RNA repertoires associated with distinct excitatory and inhibitory synapse subtypes. Pairing PSD-95–Halo-seq with cell-type–specific (such as the CaMKIIα promoter for specific excitatory neurons) or activity-dependent promoters (such as E-SARE or c-fos based promoters), ribosome profiling, proteomics, and RNA–protein interactome mapping could clarify how nanoscale RNA composition influences local translational output and synaptic function. Alternative synaptic protein targeting approaches can also yield interesting findings in similar experiments. FingR domains for CaMKIIα (Mora et al., 2013) and gephyrin (Gross et al., 2013) have been reported. FingR domains for gephyrin may reveal whether sex-dependent isoform programs extend beyond excitatory synapses. Further, applying this approach across behavioural paradigms, hormonal states, and developmental stages will be essential for determining how sex-dependent isoform programs generalize across circuits. Finally, future studies using compartment-specific manipulation of splicing or RNA localization—including CRISPR-based isoform editing or targeted perturbation of synaptic RBPs—will be critical for establishing the causal roles of identified isoforms in plasticity and memory.

### Significance

Despite these limitations, PSD-95–Halo-seq reveals previously inaccessible principles of synaptic RNA organization, demonstrating that (1) the excitatory post-synapse contains a structurally diverse and highly compartmentalized RNA repertoire, (2) sex is a major axis shaping these RNA populations, and (3) behavioural experience remodels synaptic isoform architecture in a sex-dependent manner.

These findings provide a molecular framework for understanding how synapses encode information and offer a new strategy for dissecting the compartment-specific RNA mechanisms governing learning and memory. These findings may also help explain sex-biased vulnerability to disorders involving synaptic dysfunction, including PTSD, depression, and autism.

## Supplemental Data

**Figure S1:**
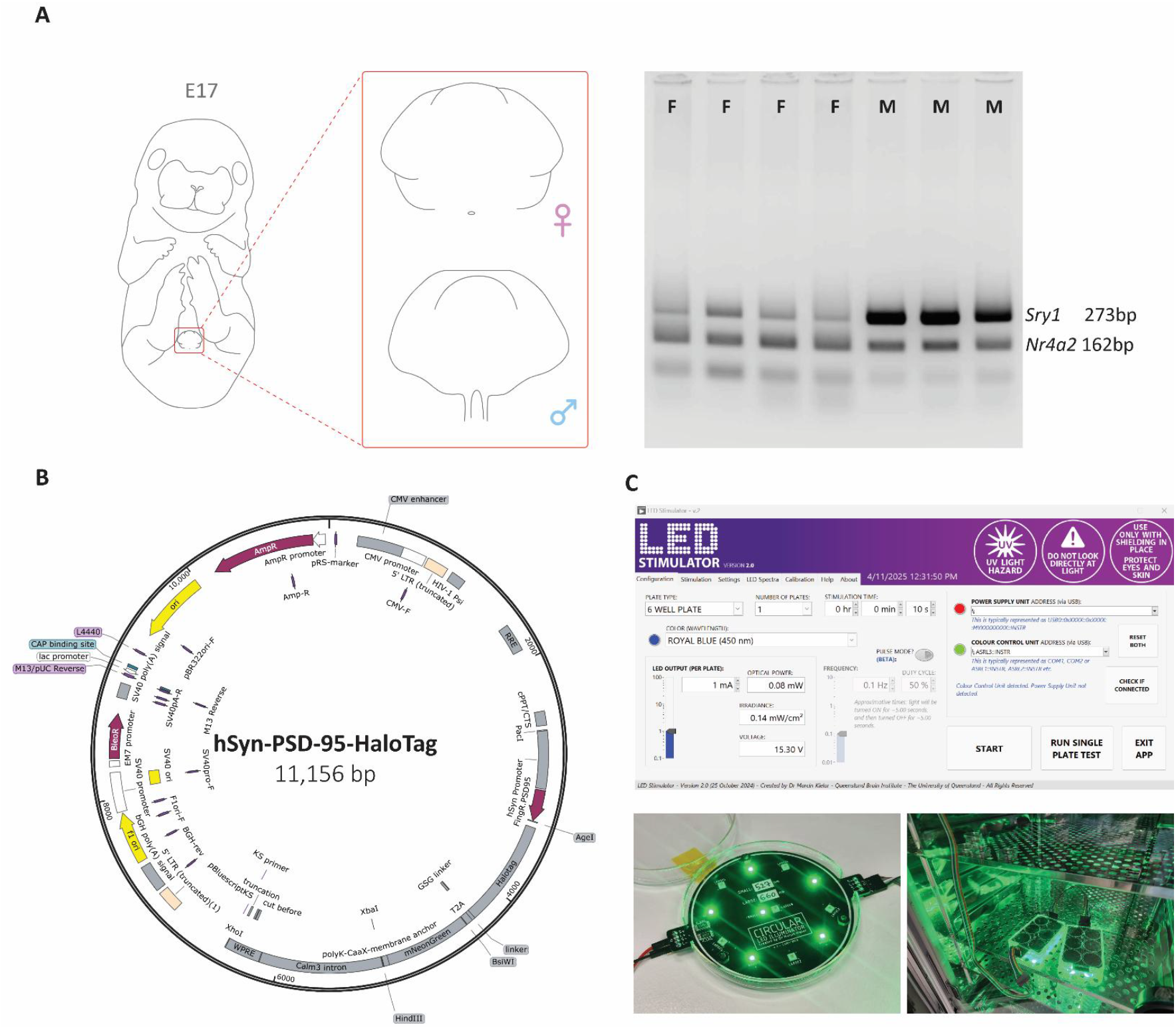
Generating and labelling sex-specific embryonic cultures with Halo-seq. **A.** Embryonic sexing technique used to derive sex-specific PCN cultures combining visual sexing of late gestational embryo genitalia (left) and validation of visual sexing with PCR of the *Sry* gene to confirm biological sex (right). **B.** Plasmid map for the hSyn-PSD-95-HaloTag construct, expressing a synapse localized HaloTag fused to FingR domain for PSD-95. **C.** Bespoke photo stimulation rig designed to allow precise titration of light for initiating labelling with hSyn-PSD-95-halo-seq. High-power LEDs (for green light: L1C1-GRN1000000000, *LUMILEDS*) were calibrated using a power and energy meter console (PM100D, *Thorlabs*) and a calibrated silicon PIN photodiode (BPX61, *OSRAM*). Customized printed circuit boards and LabVIEW scripts (*National Instruments*) were designed to provide a highly accurate and stable constant current source for the LEDs.

**Figure S2:**
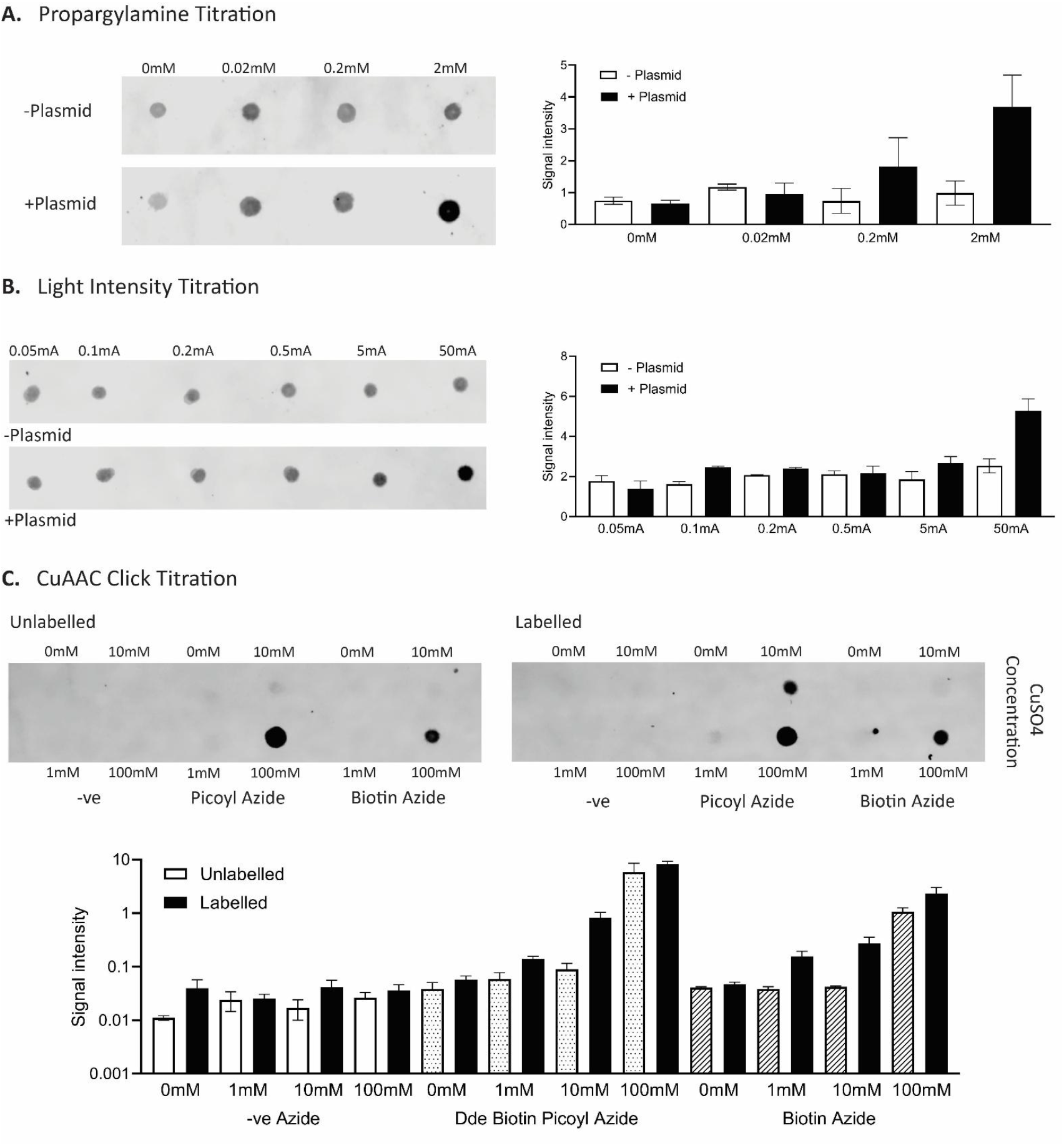
Optimization of Halo-seq parameters for labelling in Neurons. **A.** Representative dot blot visualization of biotin signal representing labelled RNA as a result of Propargylamine titration in the presences or absence of a Halo-fusion protein (+/-Plasmid) where blotted total RNA is constant (200ng). Graphed average signal intensity indicates 2mM Propargylamine as optimal for signal to noise. **B.** Representative dot blot visualization of biotin signal representing labelled RNA as a result of labelling green light intensity titration (here represented as the LED output currents from 0.05 to 50 mA) in the presences or absence of a Halo-fusion protein (+/-Plasmid) where blotted total RNA is constant (200ng). Graphed average signal intensity indicates the LED output current of 50 mA as optimal for signal to noise. For green LEDs (peak emission at 530 nm), this optimal current corresponds to a light intensity of 7.5 mW cm^-2^. We note that LEDs were current driven to enable stable and accurate light intensities – the light output of an LED is directly proportional to the forward current flowing through it (not the voltage across it). **C.** Representative dot blot visualization of biotin signal representing labelled RNA as a result of titrating copper sulphate (CuSO4) in the presence of - ve Azide, Biotin Azide or Dde Biotin Picoyl Azide. Contrasting signal in the presences or absence of labelling stimulated by irradiation with green light (Labelled/ Unlabeled) where blotted total RNA is constant (200ng). Graphed average signal intensity indicates 10mM CuSo4 with Dde Biotin Picoyl Azide as the condition with the lowest copper sulphate in which signal above noise is observed. (Group, n = 3, Error bars represent SEM)

**Table S1:** Synapse enriched RNA abundance is unaffected by sex and KCl treatment in vitro. Genes identified from PSD-95-Halo-seq samples with DESeq2 comparing synapse-enriched RNA populations. ‘All’ refers to synapse-enriched samples irrespective of KCl treatment. ‘Female’ and ‘Male’ refers to groups not treated with 50mM KCl

| DESeq2 comparison in vitro | # genes<br>pvalue<br><0.05 | # genes<br>p<sub>adj</sub><br><0.05 |
| --- | --- | --- |
| Female all vs Male all | 862 | 5 |
| Female vs Female + 50mM KCl treatment | 507 | 1 |
| Male vs Male + 50mM KCl treatment | 630 | 0 |
| Female vs Male | 773 | 1 |
| Female + 50mM KCl treatment vs Male + 50mM KCl treatment | 604 | 1 |

**Table S2:** Synapse enriched RNA abundance is affected by sex in vivo. Genes identified from PSD-95-Halo-seq samples with DESeq2 comparing synapse-enriched RNA populations. ‘All’ refers to synapse-enriched samples irrespective of behavioural training.

| DESeq2 comparison in vivo | # genes<br>pvalue<br><0.05 | # genes<br>p<sub>adj</sub><br><0.05 |
| --- | --- | --- |
| Female all vs Male all | 2093 | 1012 |
| Female Control vs Female Conditioned | 410 | 0 |
| Male Control vs Male Conditioned | 387 | 0 |
| Female Control vs Male Control | 1107 | 662 |
| Female Conditioned vs Male Conditioned | 1548 | 797 |

***Dataset 1:***Raw data for the generation of all transcriptomic data conducted in vitro.

***Dataset 2:***Raw data for the generation of all transcriptomic data conducted in vivo

